# Positive human health effects of sea spray aerosols: molecular evidence from exposed lung cell lines

**DOI:** 10.1101/397141

**Authors:** Jana Asselman, Emmanuel Van Acker, Maarten De Rijcke, Laurentijn Tilleman, Filip Van Nieuwerburgh, Jan Mees, Karel A.C. De Schamphelaere, Colin R. Janssen

## Abstract

Sea spray aerosols (SSAs) have profound effects on climate and ecosystems. Furthermore, the presence of microbiota and biogenic molecules, produced by among others marine phytoplankton, in SSAs could lead to potential human health effects. Yet the exposure and effects of SSAs on human health remain poorly studied. Here, we exposed human epithelial lung cells to different concentrations of extracts of a natural sea spray aerosol (SSA), a laboratory-generated SSA, the marine algal toxin homoyessotoxin and a chemical mTOR inhibitor. The mTOR inhibitor was included as it has been hypothesized that natural SSAs may influence the mTOR cell signaling pathway. We observed significant effects on the mTOR pathway and PCSK9 in all exposures. Based on these expression patterns, a clear dose response relationship was observed. Our results indicate a potential for positive health effects when lung cells are exposed to environmentally relevant concentrations of natural SSAs, whereas potential negative effects were observed at high levels of the laboratory SSA and the marine algal toxin. Overall, these results provide a substantial molecular evidence base for potential positive health effects of SSAs at environmentally relevant concentrations through the mTOR pathway. The results provided here suggest that SSAs contain biomolecules with significant pharmaceutical potential in targeting PCSK9.

## Introduction

Oceans and seas contain a variety of biogenic or naturally produced molecules that become airborne via sea spray aerosolization^1-3^. In addition to bacteria, which are well-known producers of biogenics, many phytoplankton species also produce a wide range of bioactive molecules such as vitamins, pigments, polyphenolics and phycotoxins^4,5^. The latter have primarily been studied in the context of harmful algal blooms, in which phycotoxins can be present at detrimental concentrations^4,6^. These toxins can through their presence in sea spray aerosols cause health effects. This has been reported for aerosolized brevetoxins which can lead to respiratory symptoms in humans during algal bloom conditions, particularly in people with asthma ^7,8^.

The effects of brevetoxins have been well-studied and documented. Little attention has, however, been given to the potential effects at low, environmentally relevant, concentrations in which phycotoxins may be present in sea spray aerosols (SSAs) during standard environmental conditions^9^. At low levels, and combined with other unidentified biogenics, these known bioactive molecules could contribute to positive health effects in coastal environments. Indeed, some of these bioactive molecules (e.g. yessotoxin^10^) have been targeted for their pharmaceutical or biotechnological potential^11,12^. Furthermore, a number of studies highlight several health promoting pathways through which airborne microbiota and biogenics from blue and green environments may have positive health effects^13,14^. Airborne microbiota are thought to contribute to a more effective immuno-regulation once inhaled or ingested^13^. Additionally, it was suggested that inhalation of low levels of microbes and parasites reduce inflammation and improve immunoregulation^13,15^. Biogenics, i.e. natural chemicals produced by among others plants, fungi, phytoplankton species and bacteria^1,3,9^, have been hypothesized to induce positive health effects via the interaction with specific cell signaling pathways (e.g. PI3K/Akt/mTORC1)^14^. The link between the mTOR pathway and positive health effects is supported by a large number of studies^16-20^ demonstrating that inhibition of this cell signaling pathway is associated with health benefits such as anti-cancer, positive cardiovascular and anti-inflammatory effects.

To date, no study has focused on the general health effects of SSAs under standard coastal conditions. Here, we aimed to explore possible molecular mechanisms that could explain health effects of SSAs at different concentrations representing low environmentally relevant concentrations as well as high potential harmful concentrations. To this end, we exposed human epithelial lung cells to extracts of (1) a natural SSA, (2) a SSA generated in a laboratory tank inoculated with homoyessotoxin-producing algae, (3) the pure bioactive molecule homoyessotoxin (hYTX) and (4) a chemical inhibitor of the mTOR pathway (Torkinib/PP242). We specifically selected hYTX and a hYTX producer as the effects of hYTX in humans remain relatively unknown despite it being structurally related to brevetoxin^10^. In addition, yessotoxin has been reported to show potential as an anti-cancer drug^10^. As such, aerosols of this phycotoxin could be a source of positive health effects. The different treatments, including different dose levels per treatment, allowed us to study a range of conditions, from most realistic, i.e. natural SSA, to the simplest, i.e. a single biogenic molecule (homoyessotoxin).

## Results & Discussion

### A small set of genes is significantly differentially expressed in all treatments

We quantified the expression of 16,5654 genes and observed differential expression across all treatments. The highest number of differentially expressed (DE) genes was observed in the pure homoyessotoxin treatment, hereafter referred to as hYTX. We observed a decreasing number of differentially expressed genes in the chemical inhibitor treatment, hereafter referred to as mTOR inhibitor, the natural SSA treatment and the treatment with a SSA generated in a laboratory tank, hereafter referred to as lab SSA. We observed almost no significant DE genes in the mid and low dose levels at false discovery rates (FDR) of 0.01 and 0.05 (Figure 1A). Given the small difference between the two FDRs, the most conservative FDR was selected for further analysis. We identified two DE genes shared by all (high dose level) treatments and the mTOR inhibitor (Figure 1B).

**Figure 1.**
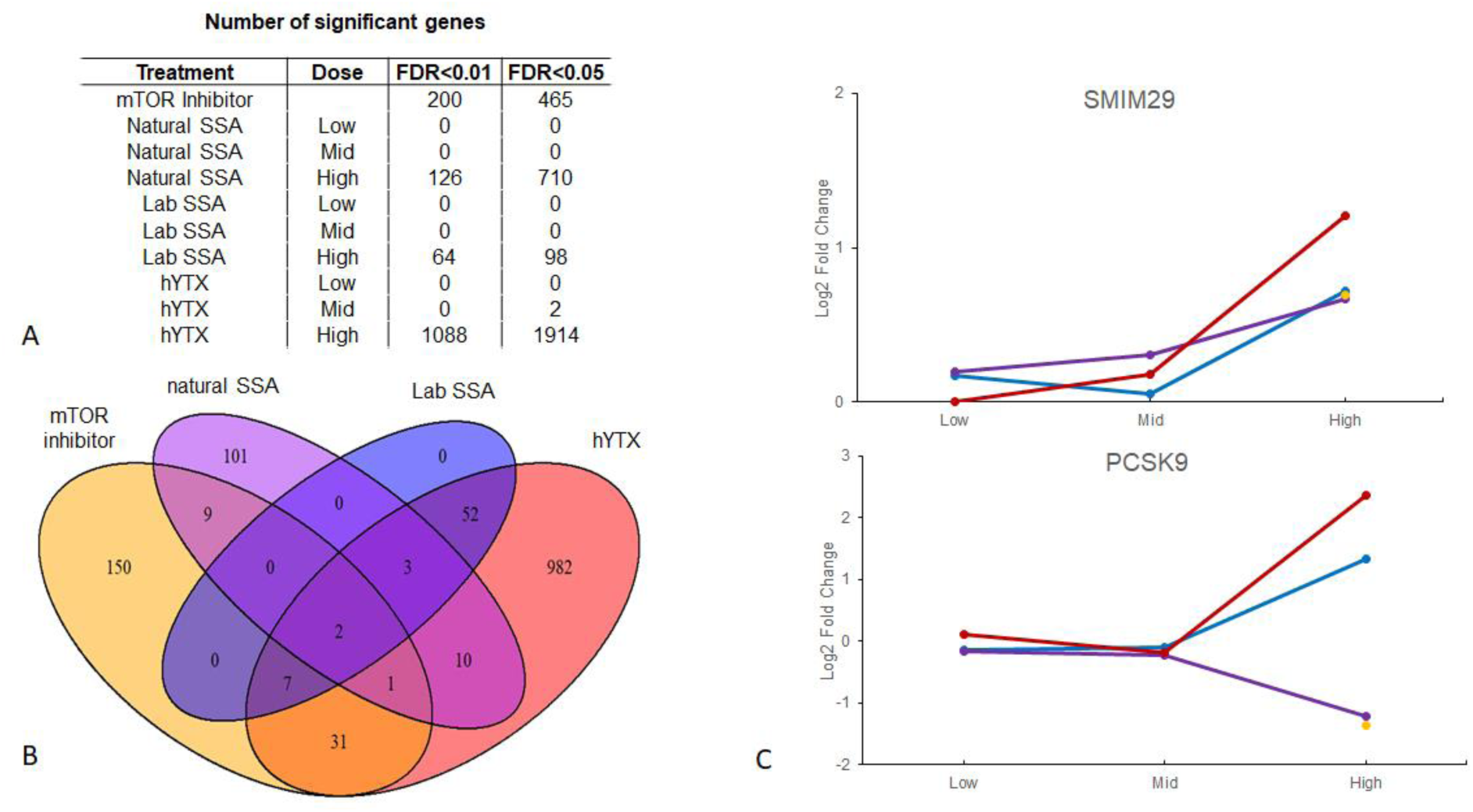
Differential gene expression across treatments. (a) Number of significant genes at different false discovery rates (FDR) for the different sea spray aerosols (SSA) treatments and homo-yessotoxin (hYTX). (b) Venn diagram of shared significant genes across treatments with significant genes at an FDR of 0.01. (c) Log fold change for small integral membrane protein 29 (SMIM29) and proprotein convertase subtilisin/kexin type 9 (PCSK9) in all treatments: natural SSA in purple, lab SSA in blue, hYTX in red and mTOR inhibitor in yellow.

The first gene was the small integral membrane protein 29 (SMIM 29). Little functional information on this protein is available, although it is ubiquitously expressed in at least 25 tissues^21^. The other gene is proprotein convertase subtilisin/kexin type 9 (PCSK9), primarily involved in lipid homeostasis and apoptosis^22^. For SMIM 29, we observed a similar pattern across all treatments with low gene expression values in low and mid dose levels, and a significant upregulation in all high dose level treatments and the mTOR inhibitor (Figure 1C). For PCSK9, the pattern is more complex. Again, we observed low gene expression values at low and mid dose levels. However, for the high dose levels, we observed a significant upregulation for hYTX and the lab SSA, while we observed a significant downregulation for the natural SSA treatment and the mTOR inhibitor (Figure 1C). For both PSCK9 and SMIM29 the effects of the lab SSA were similar but weaker than the effects of the hYTX itself. Furthermore, all DE genes that were significantly regulated by the lab SSA are a subset of the DE genes regulated by hYTX. This suggests that the effects of the lab SSA are most likely comparable to effects of a diluted hYTX treatment. Or, in other words, the effects of the lab SSA produced by a hYTX producing algae are weaker than the effects of hYTX itself despite containing the same amount of hYTX. Given that the dose levels for both treatments (lab SSA and hYTX) contain the same amount of hYTX, these results suggest that (1) lab SSAs may contain additional molecules which interact with hYTX leading to weaker effects or that (2) some of the hYTX in the lab SSA is partially degraded leading to potentially weaker effects as less “pure” hYTX is available. Literature reports only briefly on the organic composition of SSAs, but suggests a large diversity in biogenic compounds^23,24^. PCSK9 is thought to have two major functions: (1) maintenance of lipid homeostasis by the regulation of low-density lipoprotein receptors and (2) the regulation of neural apoptosis^22^. In general, the overexpression of PCSK9 is associated with the dysregulation of pathways involved in the cell cycle, inflammation and apoptosis while the inhibition of PCSK9 in carcinogenic lung cells has been associated with apoptosis of these cell lines^22^. In mouse, a similar pattern has been observed^25^. Overexpression of PCSK9 was associated with multi-organ pathology and inflammation while PCSK9 deficiency was associated with protection against inflammation, organ pathology and systemic bacterial dissemation^25^. Based on these findings in literature and the similarities between the PCSK9 expression patterns of the mTOR inhibitor and the natural SSA, our results suggest a potential for positive health effects when lung cell lines are exposed to natural SSA. Based on the results provided here on PCSK9, we propose that SSA contain molecules with significant pharmaceutical potential in targeting PCSK9^26^.

### Significant effects on the mTOR regulatory pathways differ across treatments

The biogenics hypothesis suggests that the mTOR pathway is one of the key drivers of the coastal induced health benefit. Here both the KEGG mTOR pathway annotation^27^ and the molecular signature databases^28^, which contains a hallmark set of genes upregulated upon activation of the mTORC1 complex, were used to test this hypothesis. No significant enrichment of the KEGG mTOR pathway in any of the treatments was observed. Individual genes of the mTOR pathway, however, were significantly regulated in different high dose treatments, with the exception of the lab SSA for which no mTOR genes were differentially expressed (Table S1). We also noted significant enrichment scores of the GSEA Hallmark mTORC1 set for all high dose treatments, excluding the natural SSA, and the mTOR inhibitor (Table S2). Taking a closer look at the hallmark mTORC1 set, we concluded that the gene expression patterns differed across treatments (Figure S1). Hierarchical clustering of these patterns indicated that DE genes were in general regulated in the opposite direction for hYTX and the lab SSA versus the natural SSA and the chemical inhibitor (Figure S1). This pattern is even more prominent when focusing on the genes that contribute significantly to the enrichment score in the hallmark set for all 4 treatments (Figure 2).

This group of 17 genes showed completely opposite regulation patterns in the high dose hYTX versus the high dose natural SSA and the chemical inhibitor (Figure 2). The high dose laboratory SSA showed a similar but less intense and weaker regulation than the high dose pure hYTX. Overall, these results suggest that all treatments affect the mTOR pathway but the effects and the potential positive health effects differ across treatments. Interestingly, the effects of the natural SSA closely resemble the effects of the mTOR inhibitor, but contrast with the effects of hYTX and the lab SSA. In addition, we again observed that the lab SSA caused effects in a similar direction as the pure hYTX, albeit weaker. The differences between the natural SSA on one hand and the pure hYTX and lab SSA on the other hand highlight that while all treatments target the mTOR pathway, their effects are opposite. This may suggest (1) that the natural SSA contains different molecules than the lab SSA and hYTX or (2) that less “pure” hYTX is available due to degradation of hYTX. Both assumptions suggest a lower bioavailability of pure hYTX, potentially leading to a lower actual dose. This is also supported by the observation that six genes of the mTOR pathway show a significant dose response effect (Table S2). The similarities in regulation of the mTOR pathway between the natural SSA and chemical inhibitor suggest that natural SSAs contain molecules that cause similar effects on the mTOR pathway as the chemical inhibitor.

**Figure 2.**
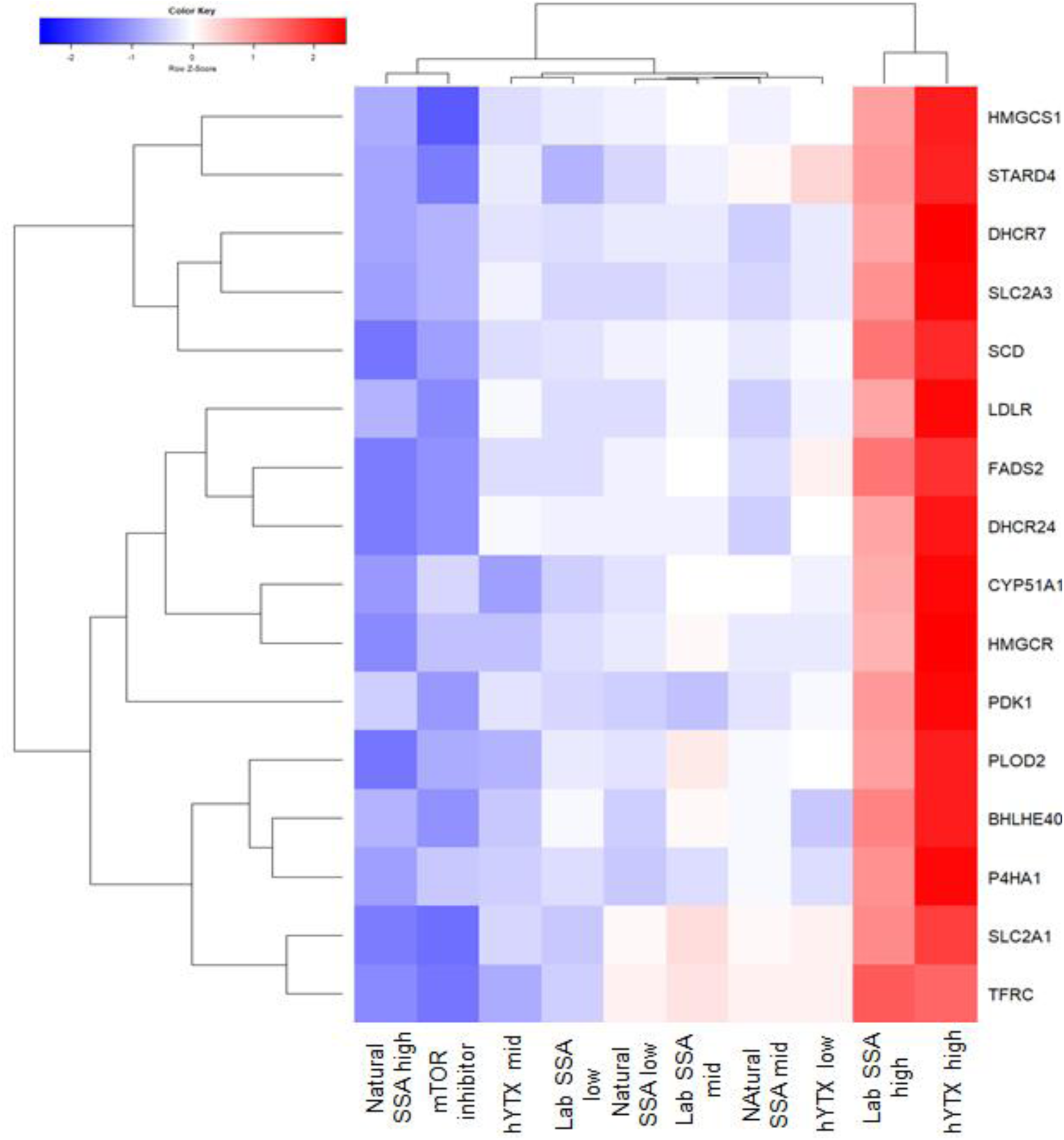
Enrichment of the mTOR Hallmark set. (a) Heatmap for all treatments of the fold changes of genes that contribute significantly to the enrichment score for all three treatments at the highest dose and the mTOR inhibitor. Treatments: chemical inhibitor, homo-yessotoxin (hYTX), lab sea spray aerosol (SSA) and natural sea spray aerosol (SSA) at low, mid and high doses.

### Significant concentration response patterns across treatments

We observed a total of 1898 genes with a significant dose response effect across the three treatments (hYTX, lab SSA and natural SSA). Based on a regression analysis and clustering, we found four clusters of dose response patterns. These clusters all show the same trend which consists of a steep dose response curve for hYTX while the lab SSA and the natural SSA show a slower increase (Figure S2). When observing gene expression patterns for the clusters across all treatments, we see the same pattern of two groups as reported in the sections above, one containing the high dose hYTX and lab SSA treatment while the other contains the remaining treatments. In three of the four clusters, the mTOR inhibitor treatment clustered together with the high dose natural SSA treatment. These clustering results suggest that the observations we have made above for the mTOR pathway and PCSK9 gene are not limited to these two observations but can be extended to all genes with a significant dose response effect. A pathway analysis highlighted four pathways that were enriched for genes with a significant dose response effect (Table S3). These pathways are the spliceosome, lysosome, steroid biosynthesis and glycogenesis. For all these pathways, we observed two major clusters (Figure 3A-D, Figures S3-6): the pattern for the highest dose hYTX was again similar to that of the highest dose lab SSA while the pattern of the natural SSA was again similar to that of the mTOR inhibitor. Again, we observed the opposite regulation of genes in these two groups for three pathways. Indeed, for the lysosome, steroid biosynthesis and glycogenesis pathways we noted an upregulation of genes with a significant dose-response effect in the high dose hYTX and the high dose lab SSA (Figure 3B-D). In contrast, for these same pathways, we observed primarily a downregulation of the same significant genes in the mid and low dose hYTX and lab SSA, as well as all natural SSA treatments and the chemical inhibitor with the exception of the low lab SSA in the glycogenis pathway. For the spliceosome, we observed significant upregulation in all treatments with exception of the low and mid hYTX and mid lab SSA (Figure 3A).

**Figure 3.**
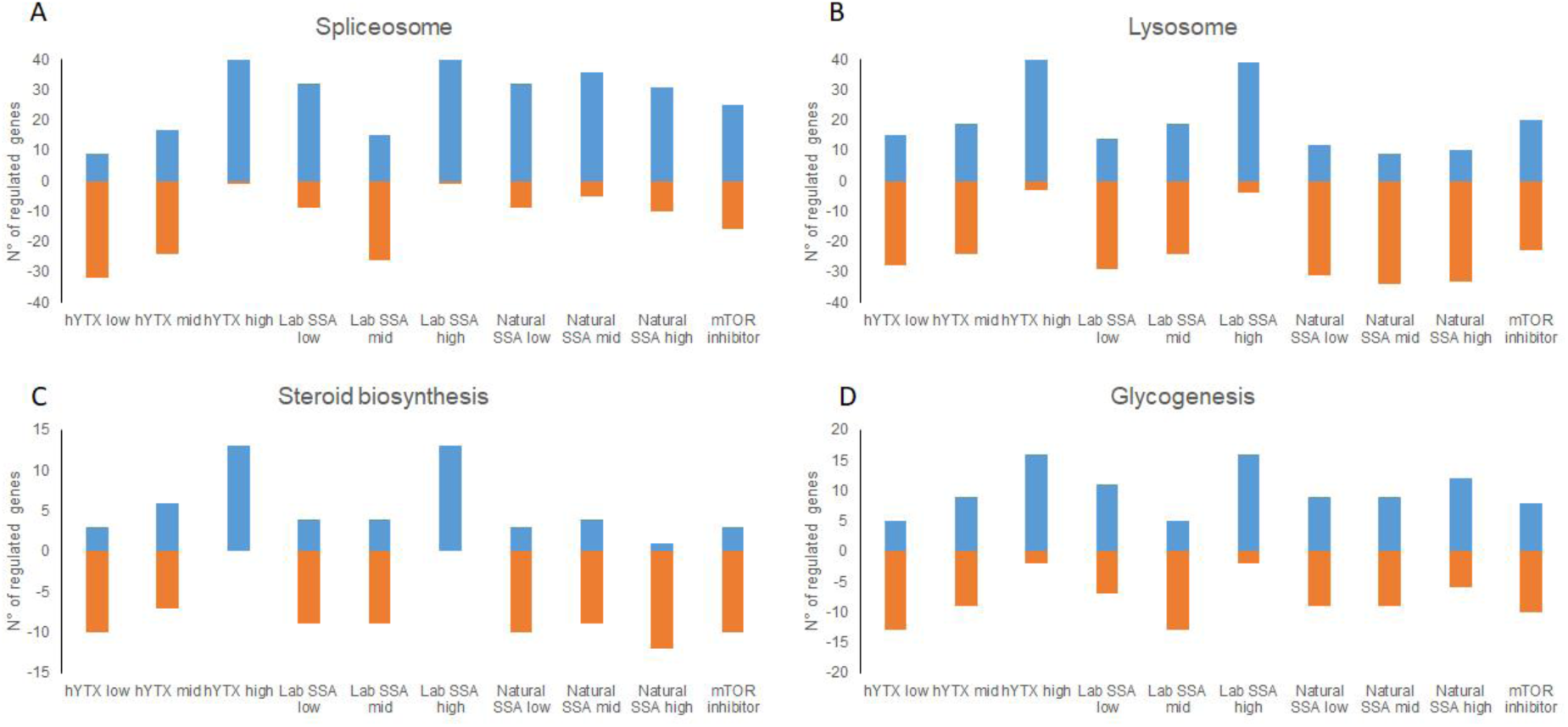
Dose response patterns in significant pathways. Number of significantly upregulated (>0) or downregulated (<0) genes in **(A)** the spliceosome, **(B)** the lysosome, **(C)** steroid biosynthesis, **(D**) glycogenesis for all treatments: natural sea spray aerosols (SSA) lab sea spray aerosol (SSA), homoyessotoxin (hYTX) and mTOR inhibitor.

Similar to the effects in the mTOR pathway, the effects on these pathways again show a similar regulation of genes for the natural SSA and the mTOR inhibitor. For the steroid biosynthesis, these results are not surprising given the links that have already been discussed above between mTOR and lipid biosynthesis. In addition to steroid biosynthesis, the lysosome and glycogenesis also have links to mTOR. The inhibition of the mTOR pathway is known to activate protein degradation and autophagy through among others the lysosome^29,30^. The spliceosome has been proposed as a therapeutic target in cancer cells to inhibit mTOR, which led to autophagy^31^. Specifically, depletion of small nuclear ribonucleoprotein polypeptide E (SNRPE) led to reduced cell viability in lung cancer cell lines. Here, we observed in addition to dose response effects for the spliceosome, also a significant downregulation of SNRPE in the highest hYTX treatment but not in any of the other treatments (Table S4). Overall, the pathways with significant dose response effects can all be indirectly linked to the mTOR pathway, suggesting that the effects here are a consequence of the effects on the mTOR pathway, which most likely induce a cascade of events and interactions with other pathways.

### Significant effects unique to hYTX and sea spray aerosols

While we have focused on similarities between effects of our experimental treatments and the mTOR inhibitor, we also observed effects unique to these treatments. We observed the differential expression of three genes shared by all high dose treatments. These genes are stearoyl-CoA desaturase (SCD), cytochrome P450 family 1 subfamily B member 1 (CYP1B1) and peptidyl arginine deiminase 3 (PADI3). For SCD, we observed a pattern similar to that of the PCSK9 expression, i.e. exposure to the natural SSA led to downregulation while exposure to the lab SSA and hYTX led to upregulation (Figure 4A). This can be attributed to the functions of these genes (i.e. SCD, PCSK9), as both are involved in lipid biosynthesis. Furthermore, research has already indicated links between the mTOR pathway and the lipid homeostasis^32^, including the effects on SCD and other genes after exposure to mTOR inhibitors^32^. Evidence points to sterol regulatory element binding transcription factor 1 (SREBF1) through which the regulation of lipogenesis by mTOR is achieved^32^. This gene was significantly regulated by the natural SSA, but not by any of the other treatments.

**Figure 4.**
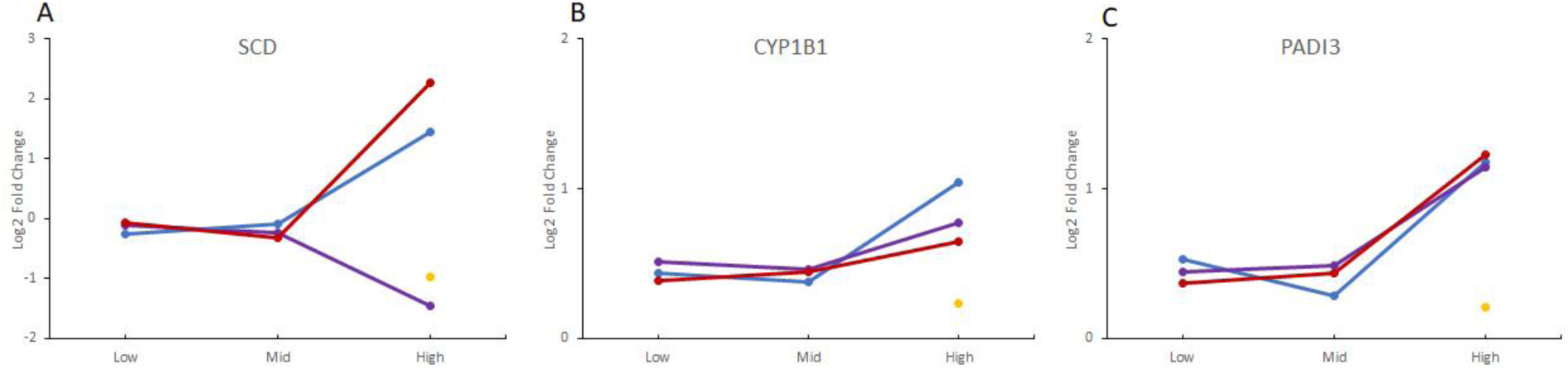
Differential gene expression in hYTX and sea spray aerosol treatments. Log fold change for **(A)** stearoyl-CoA desaturase (SCD), **(B)** cytochrome P450 family 1 subfamily B member 1 (CYP1B1) and **(C)** peptidyl arginine deiminase 3 (PADI3) for all treatments: natural sea spray aerosol (SSA) in purple, lab sea spray aerosol (SSA) in blue, homoyessotoxin (hYTX) in red and mTOR inhibitor in yellow.

For CYP1B1 and PADI3, we observed a pattern similar as for SMIM29, in which all treatments resulted in significant upregulation (Figure 4B-C). The first is commonly involved in the metabolism of xenobiotics and could play a role in metabolizing some of the biogenic molecules. Literature has also reported a relation between CYP1B1 and SCD in lipid homeostasis in liver cells^33^, although the extent of this relation in lung cells remains unclear. Overexpression of CYP1B1 has also been reported in lung cell lines through the aryl hydrocarbon receptor^34^, but no significant effects for this receptor were observed in any treatment of our study(Table S4). This suggests that the overexpression of CYP1B1 is more likely related to the regulation of SCD. Lastly, we observed a significant upregulation of PADI3 in all three high dose treatments (hYTX, lab SSA and natural SSA). PADI3 is generally not expressed in lung cells^21^ and is primarily expressed in epidermis cells and keratinocytes^35^. Its function in lung cell lines remains unclear. Overall, we observe here differential expression of genes linked to the mTOR pathway in all three high dose treatments (natural SSA, lab SSA, and hYTX). Most likely, the effects on these genes are caused by the primary effects on the mTOR pathway. Furthermore, these effects while linked to the mTOR pathway, are not observed with the mTOR inhibitor. This suggests that the effects of these experimental treatments (natural SSA, lab SSA, and hYTX) extend beyond the inhibition of mTOR but are related to or initiated by the effects on the mTOR pathway.

### Differences in dose level lead to a different regulation of the same significant genes and pathways across treatments

The results of gene set and pathway enrichments as well as individual genes highlight that the effects are primarily mediated or linked through the mTOR pathway (Figure 5). Here, we studied both a natural and a lab SSA, as well as the effects of pure hYTX as potential key biogenic molecule in natural SSAs. While we observed similar pathways, and to some extent, similarly affected genes across these different treatments, the regulation was not necessarily the same. The hYTX and the lab SSA showed a similar pattern across all pathways and genes, while differences were observed with the natural SSA and the chemical inhibitor. These differences could be related to the differences in doses. The high dose treatment for both hYTX and lab SSA of 0.5 µg liter^-1^ is an extreme case scenario, reflecting concentrations in water during harmful algal blooms (supportive information 1.2). The environmental (background) concentrations of hYTX in water and air have not been previously reported but are expected to lie between the low and mid dose levels based on estimates of cell counts of hYTX producers and hYTX production per cell (supportive information 1.2). As such, it is clear that there is a switch in effects where at high doses hYTX and lab SSAs can cause negative effects while the regulation of pathways and genes is the opposite at low and mid doses, suggesting positive health effects at environmentally relevant (background) concentrations. A direct comparison can only be made with the lab SSA in terms of total aerosols by using the cation sodium as a proxy for aerosolization^36^. We observed that the lab SSA dose levels are 2.8 µg Na^+^ well^-1^, 0.06 µg Na^+^ well^-1^ and 0.00006 µg Na^+^ well^-1^ while the natural SSA dose levels, due to the smaller sample size, were 0.6 µg Na^+^ well^-1^, 0.14 µg Na^+^ well^-1^ and 0.014 µg Na^+^ well^-1^ (section 1.2). As such, the highest dose for the natural SSA contains only 20% of the amount of aerosols in the high dose lab SSA treatment. This supports the assumptions made above that exposure to environmentally relevant concentrations of marine biogenics, sampled from the environment, can lead to positive health effects at environmentally relevant concentrations. In addition, we observed similar patterns of gene expression for the mTOR inhibitor and the highest natural SSA treatment.

**Figure 5.**
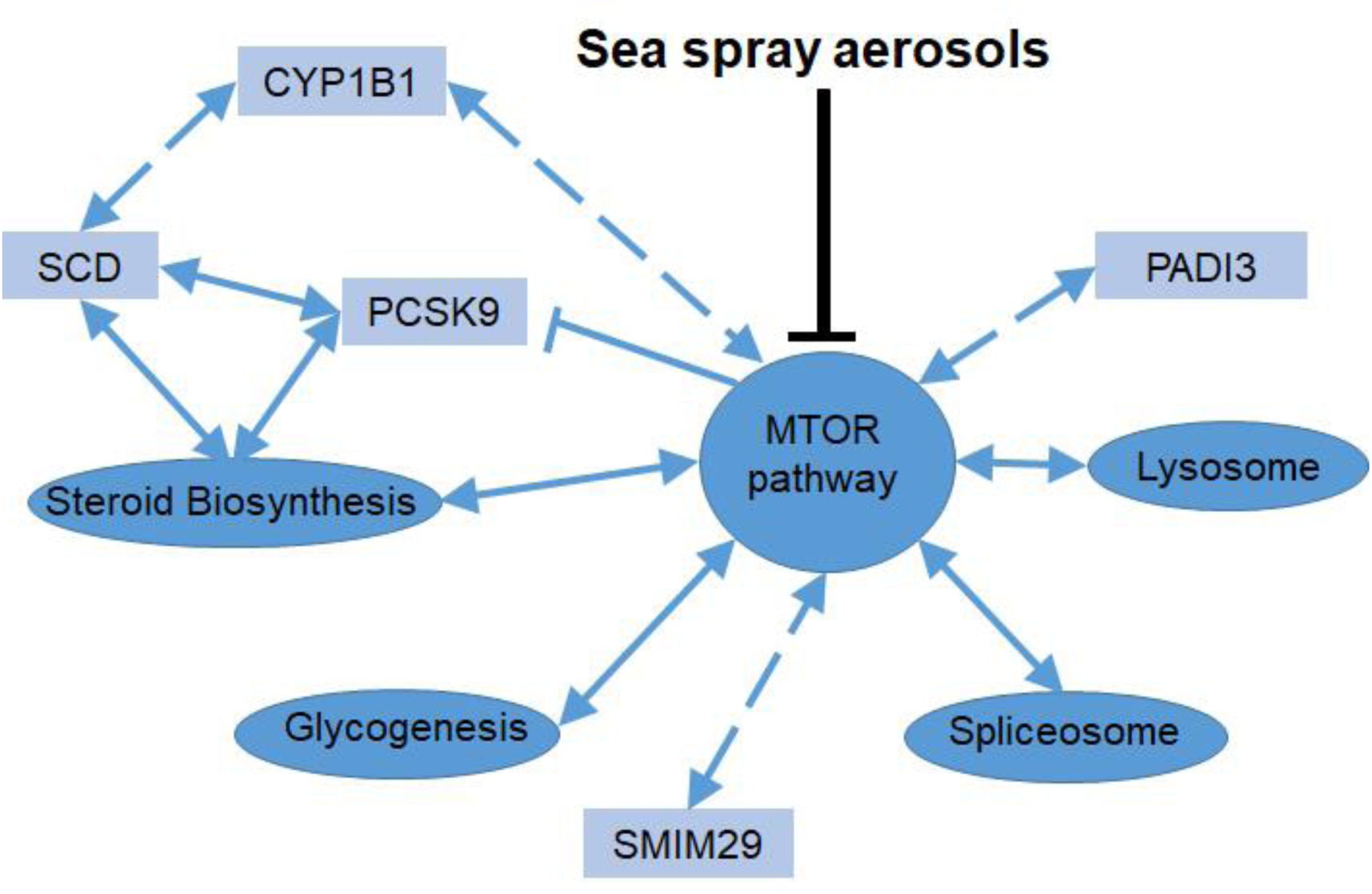
Molecular effects of marine aerosolized biogenics. A schematic representation of the molecular effects of sea spray aerosols observed within this study. Pathways are represented by ellipses, genes are represented by rectangles. Solid blue arrows represent interactions with a solid evidence base, dashed arrows represent hypothetical interactions observed, ⊢ represent inhibition.

Overall, our results support the biogenesis hypothesis postulated by Moore^14^ that marine airborne biogenics interact with the mTOR pathway potentially leading to health benefits. We report significant effects on the mTOR pathway in all treatments, though differences in regulation of this pathway were observed. Furthermore, significant genes and enriched pathways across treatments all interact with mTOR, indicating that marine biogenics trigger a cascade of events through interaction with the mTOR pathway (Figure 5).Thus, the effects of marine airborne biogenics are not limited to the mTOR pathway but include a cascade of genes and pathways involved in different metabolic processes (e.g. steroid biosynthesis, lysosome) with key links to mTOR (Figure).

## Methods

### Culturing of A549 cells

Adenocarcinoma alveolar basal cell lines (A549) were maintained in Dulbecco’s Modified Eagle Medium (DMEM), supplemented with 10% fetal bovine serum and 5000 units.mL^-1^ penicillin-streptomycin at 37°C, 5% CO_2_ and >95% relative humidity. Confluent cell cultures (after 2-3 days) were passaged via trypsination (0.5% trypsin-EDTA) and split in a ratio 1:6.

### Experimental procedure

Confluent cell cultures were trypsinized and transferred in 3mL fresh DMEM to Nunc 6-well multiplates at a density of 320,000 cells.well^-1^. After seeding, cells were incubated for 10 hours at 37°C, 5% CO2 and >95% relative humidity to stimulate growth and adherence to the surface. Then, cells were subjected to one of five treatments: (1) negative control, (2) an extract of a natural SSA sample from the seashore, (3) an extract of a laboratory generated SSA, (4) homoyessotoxin, (5) a chemical inhibitor of the mTOR pathway, i.e. Torkinib or PP242 (LC Laboratories). The multiwell plates were then incubated for another 43 hours at identical conditions prior to RNA extraction. The negative control treatment also contained 2% methanol to exclude a solvent effect as all other treatments were extracted, diluted or dissolved in methanol. The chemical inhibitor treatment consisted of 0.3 µM of Torkinib or PP242. The natural sea spray aerosol sample was collected on a Whatman QM-A Quartz Microfiber filter at the waterline close to Ostend, Belgium (51°14’27”N, 2°56’10”E) by sampling for 46 minutes at a flow of 10 L min^-1^, which corresponds to the minute ventilation of an average human in rest (9-10 L min^-1^)^37,38^. During sampling, the wind direction was 0.7 ± 3.1 ° (North), speed was 15.0 ± 0.6 m s^-1^, indicating white cap SSA production. The detailed sampling and extraction procedure is described in supportive information 1.1. The lab SSA was obtained by inoculating a marine aerosol reference tank^39^ with 10^6^ cells L^-1^ of *Protoceratium reticulatum*, a hYTX producer (SCCAP K-1474), and collecting the generated SSA on a Whatman QM-A Quartz Microfiber filter at a flow of 10 L min^-1^ for 16 hours to obtain sufficient material for further experiments and analysis. The detailed procedure is described in supportive information 1.1. Filters of the natural SSA and lab SSA were extracted following the same methanol extraction procedure. Certified reference material of hYTX was commercially obtained (National Research Council Canada) as a liquid with a concentration of 5µM hYTX dissolved in methanol. This reference material was further diluted in methanol to obtain the following dose levels: 0.5 µg L^-1^ (high), 0.01 µg L^-1^ (mid), 0.00001 µg L^-1^ (low). Concentrations of hYTX in the lab SSA were measured using ultra-high-performance liquid chromatography high-resolution Orbitrap mass spectrometry following procedures as reported by Orellana et al. (2014)^40^. To allow an optimal comparison between the hYTX treatment and the lab SSA, the lab SSA dose levels were determined based on the measured hYTX in these samples and the same dose levels as the hYTX treatment were selected (0.5 µg L^-1^ (high), 0.01 µg L^-1^ (mid), 0.00001 µg L^-1^ (low)). For natural SSA, low, mid and high doses were determined by comparing the total alveolar surface with the cell surface available in a single well (9.6 cm^2^) and comparing the sample collection duration (46 min) and experimental exposure duration (43 h), see supportive information 1.2. We selected a low dose that represents the same exposure as the amount of inhaled SSA during the sampling period at the seashore but extended over an 43 h exposure period and normalized to the cell surface in a single well (detailed calculations are reported in supportive information, section 1.2). The mid and high dose represent a 10x and 40x concentration of the low dose level. These levels were specifically chosen to adhere to environmentally realistic (background) concentrations. The mid dose level (10x concentration) was based on the hypothesis of increased minute ventilation during physical exercise which is reported to vary between 70-100 L min^-1^ for both continuous and intermittent exercise^38,41,42^. The high dose level (40x concentration) was selected based on the hypothesis of increased aerosolization (i.e. improved wind conditions) as well as activities at the shore line or at sea (e.g. swimming, sailing, windsurfing,..,..). Detailed procedure is described in the supportive information, section 1.2.

### RNA extraction, library preparation and sequencing

RNA was extracted using the Qiagen RNEasy kit following the manufacturer’s instructions including DNAse digestion. After RNA extraction, the concentration and quality of the total extracted RNA was checked by using the ‘Quant-it ribogreen RNA assay’ (Life Technologies, Grand Island, NY, USA) and the RNA 6000 nano chip (Agilent Technologies, Santa Clara, CA, USA), respectively. Subsequently, 250 ng of RNA was used to perform an Illumina sequencing library preparation using the QuantSeq 3’ mRNA-Seq Library Prep Kits (Lexogen, Vienna, Austria) according to manufacturer’s protocol. During library preparation 14 PCR cycles were used. Libraries were quantified by qPCR, according to Illumina’s protocol ‘Sequencing Library qPCR Quantification protocol guide’, version February 2011. A High sensitivity DNA chip (Agilent Technologies, Santa Clara, CA, US) was used to control the library’s size distribution and quality. Sequencing was performed on a high throughput Illumina NextSeq 500 flow cell generating 75 bp single reads.

### Data analysis

Per sample, on average 7.5 x 10^6^ ± 1.6 x 10^6^ reads were generated. First, these reads were trimmed using cutadapt^43^ version 1.15 to remove the “QuantSEQ FWD” adaptor sequence. The trimmed reads were mapped against the Homo sapiens GRCh38.89 reference genome using STAR^44^ version 2.5.3a. The RSEM^45^ software, version 1.3.0, was used to generated the count tables. Differential gene expression analysis between groups of samples was performed using edgeR^46^. Genes with less than 1 cpm in less than 4 samples were discarded, resulting in 16,546 quantifiable genes. Read counts were normalized using trimmed mean of M-values (TMM) followed by a pairwise comparison of treatments with the negative and positive control using an exact test^46^. Significantly differentially expressed (DE) genes were called at a false discovery rate of 0.01. Significant enrichment of KEGG pathways^27^ with DE genes was done using a fisher test and called at an adjusted p-value level of 0.01. Benjamini-Hochberg adjustment was used to account for multiple testing. Gene set enrichment analysis (GSEA) was conducted to detect enrichment in hallmark gene sets and genetic and chemical perturbations gene sets of the molecular signature database^28^. Enriched gene sets were identified at a false discovery rate of 0.01. A dose response analysis was performed with the maSigPro^47^ R package for each of the three treatments of algal toxins. In a first step a general linear model was built with the 3 treatments, the 3 concentrations and the square of each of the 3 concentrations. Statistical testing was done using the log-likelihood ratio statistic. Genes with a FDR < 0.05 were considered significantly differential. In a second step, for each significant differentially expressed gene, an optimized regression model was created using stepwise backward regression. Exclusion of the quadratic term from the model was performed using a regression ANOVA, testing if the regression coefficients differ from 0 at a significance level of 0.05. Afterwards the goodness of fit, R^2^, of each optimized regression model was computed. Genes with a goodness of fit greater than 0.8 were used in a hierarchical cluster analysis based on the correlation between the regression models of the genes.

## Data availability

Raw and processed sequencing reads are deposited in GEO and available under accession number: <u>GSE113144</u>.

## Acknowledgements

The authors acknowledge Jolien Depecker, Emmy Pequeur, Illias Semmouri for their assistance during the experiment, Nancy De Saeyer, Marc Van Der Borght, Michiel Suerinck, Sam Baelus and the staff of nxtgnt for their technical assistance. The cell lines have been originally kindly donated by Ilse Beck and Marc Bracke from the Laboratory of Experimental Cancer Research (UGent). The LC-MS analysis was conducted by using the equipment provided by Lynn Vanhaecke and Steve Huysman from the Laboratory for Chemical Analysis (UGent). This research was supported by a Brilliant Marine Research Grant awarded by VLIZ Philantrophy to Emmanuel Van Acker. Jana Asselman is a postdoctoral fellow of the Science Foundation Flanders (FWO). The authors declare that they have no competing financial interests.

## Author contributions

EVA, MDR and CJ conceptualized the idea and research question. EVA directed the sea spray aerosol sampling, production and extraction with input from MDR. EVA designed and executed the experiment with input from JA, KDS and CJ. FVN and JA developed the sequencing design. JA and LT processed and analyzed the data. JA wrote the manuscript with input from EVA. MDR, JM, FVN, KDS and CJ reviewed and edited the manuscript.

## References

1 Prather, K. A. et al. Bringing the ocean into the laboratory to probe the chemical complexity of sea spray aerosol. P Natl Acad Sci USA 110, 7550-7555, doi:10.1073/pnas.1300262110 (2013).

2 Leck, C. & Bigg, E. K. Biogenic particles in the surface microlayer and overlaying atmosphere in the central Arctic Ocean during summer. Tellus B 57, 305-316, doi:DOI 10.1111/j.1600-0889.2005.00148.x (2005).

3 Leck, C. & Bigg, E. K. Source and evolution of the marine aerosol - A new perspective. Geophys Res Lett 32, doi:Artn L1980310.1029/2005gl023651 (2005).

4 Van Dolah, F. M. Marine algal toxins: Origins, health effects, and their increased occurrence. Environ Health Persp 108, 133-141, doi:DOI 10.1289/ehp.00108s1133 (2000).

5 de Morais, M. G., Vaz Bda, S., de Morais, E. G. & Costa, J. A. Biologically Active Metabolites Synthesized by Microalgae. Biomed Res Int 2015, 835761, doi:10.1155/2015/835761 (2015).

6 Cheng, Y. S. et al. Characterization of marine aerosol for assessment of human exposure to brevetoxins. Environ Health Persp 113, 638-643, doi:10.1289/ehp.7496 (2005).

7 Fleming, L. E. et al. Exposure and effect assessment of aerosolized red tide toxins (brevetoxins) and asthma. Environ Health Perspect 117, 1095-1100, doi:10.1289/ehp.0900673 (2009).

8 Fleming, L. E. et al. Initial evaluation of the effects of aerosolized Florida red tide toxins (brevetoxins) in persons with asthma. Environ Health Perspect 113, 650–657 (2005).

9 Despres, V. R. et al. Primary biological aerosol particles in the atmosphere: a review. Tellus B 64, doi:ARTN 1559810.3402/tellusb.v64i0.15598 (2012).

10 Alfonso, A., Vieytes, M. R. & Botana, L. M. Yessotoxin, a Promising Therapeutic Tool. Mar Drugs 14, doi:10.3390/md14020030 (2016).

11 Imhoff, J. F., Labes, A. & Wiese, J. Bio-mining the microbial treasures of the ocean: new natural products. Biotechnol Adv 29, 468-482, doi:10.1016/j.biotechadv.2011.03.001 (2011).

12 Fenical, W. New pharmaceuticals from marine organisms. Trends Biotechnol 15, 339-341, doi:10.1016/S0167-7799(97)01081-0 (1997).

13 Rook, G. A. Regulation of the immune system by biodiversity from the natural environment: An ecosystem service essential to health. P Natl Acad Sci USA 110, 18360-18367, doi:10.1073/pnas.1313731110 (2013).

14 Moore, M. N. Do airborne biogenic chemicals interact with the PI3K/Akt/mTOR cell signalling pathway to benefit human health and wellbeing in rural and coastal environments? Environ Res 140, 65-75, doi:10.1016/j.envres.2015.03.015 (2015).

15 Lowry, C. A. et al. The Microbiota, Immunoregulation, and Mental Health: Implications for Public Health. Curr Environ Health Rep 3, 270-286, doi:10.1007/s40572-016-0100-5 (2016).

16 Lamming, D. W. & Sabatini, D. M. A Central Role for mTOR in Lipid Homeostasis. Cell Metab 18, 465-469, doi:10.1016/j.cmet.2013.08.002 (2013).

17 Lamming, D. W., Ye, L., Sabatini, D. M. & Baur, J. A. Rapalogs and mTOR inhibitors as anti-aging therapeutics. J Clin Invest 123, 980-989, doi:10.1172/Jci64099 (2013).

18 Sabatini, D. M. Role of the mTOR signaling pathway in disease. Inflamm Bowel Dis 12, S9-S9, doi:Doi 10.1097/00054725-200604002-00019 (2006).

19 Sabatini, D. M. mTOR and cancer: insights into a complex relationship. Nat Rev Cancer 6, 729- 734, doi:10.1038/nrc1974 (2006).

20 Corradetti, M. N. & Guan, K. L. Upstream of the mammalian target of rapamycin: do all roads pass through mTOR? Oncogene 25, 6347-6360, doi:10.1038/sj.onc.1209885 (2006).

21 Fagerberg, L. et al. Analysis of the human tissue-specific expression by genome-wide integration of transcriptomics and antibody-based proteomics. Mol Cell Proteomics 13, 397- 406, doi:10.1074/mcp.M113.035600 (2014).

22 Xu, X. H. et al. PCSK9 regulates apoptosis in human lung adenocarcinoma A549 cells via endoplasmic reticulum stress and mitochondrial signaling pathways. Exp Ther Med 13, 1993- 1999, doi:10.3892/etm.2017.4218 (2017).

23 Fu, P. Q., Kawamura, K., Chen, J., Charriere, B. & Sempere, R. Organic molecular composition of marine aerosols over the Arctic Ocean in summer: contributions of primary emission and secondary aerosol formation. Biogeosciences 10, 653-667, doi:10.5194/bg-10-653-2013 (2013).

24 O’Dowd, C. D. et al. Biogenically driven organic contribution to marine aerosol. Nature 431, 676-680, doi:10.1038/nature02959 (2004).

25 Dwivedi, D. J. et al. Differential Expression of Pcsk9 Modulates Infection, Inflammation, and Coagulation in a Murine Model of Sepsis. Shock 46, 672-680, doi:10.1097/Shk.0000000000000682 (2016).

26 Hazen, S. L. New lipid and lipoprotein targets for the treatment of cardiometabolic diseases. J Lipid Res 53, 1719-1721, doi:10.1194/jlr.E030205 (2012).

27 Kanehisa, M. & Goto, S. KEGG: Kyoto Encyclopedia of Genes and Genomes. Nucleic Acids Research 28, 27-30, doi:DOI 10.1093/nar/28.1.27 (2000).

28 Subramanian, A. et al. Gene set enrichment analysis: A knowledge-based approach for interpreting genome-wide expression profiles. P Natl Acad Sci USA 102, 15545-15550, doi:10.1073/pnas.0506580102 (2005).

29 Settembre, C. et al. A lysosome-to-nucleus signalling mechanism senses and regulates the lysosome via mTOR and TFEB. EMBO J 31, 1095-1108, doi:10.1038/emboj.2012.32 (2012).

30 Zhao, J. H., Zhai, B., Gygi, S. P. & Goldberg, A. L. mTOR inhibition activates overall protein degradation by the ubiquitin proteasome system as well as by autophagy. P Natl Acad Sci USA 112, 15790-15797, doi:10.1073/pnas.1521919112 (2015).

31 Quidville, V. et al. Targeting the Deregulated Spliceosome Core Machinery in Cancer Cells Triggers mTOR Blockade and Autophagy. Cancer Res 73, 2247-2258, doi:10.1158/0008- 5472.Can-12-2501 (2013).

32 Laplante, M. & Sabatini, D. M. An Emerging Role of mTOR in Lipid Biosynthesis. Current Biology 19, R1046-R1052, doi:10.1016/j.cub.2009.09.058 (2009).

33 Li, F. et al. Lipidomics Reveals a Link between CYP1B1 and SCD1 in Promoting Obesity. J Proteome Res 13, 2679-2687, doi:10.1021/pr500145n (2014).

34 Chang, J. H. T., Chang, H., Chen, P. H., Lin, S. L. & Lin, P. P. Requirement of aryl hydrocarbon receptor overexpression for CYP1B1 up-regulation and cell growth in human lung adenocarcinomas. Clin Cancer Res 13, 38-45, doi:10.1158/1078-0432.Ccr-06-1166 (2007).

35 Dong, S. et al. NF-Y and Sp1/Sp3 are involved in the transcriptional regulation of the peptidylarginine deiminase type III gene (PADI3) in human keratinocytes. Biochem J 397, 449- 459, doi:10.1042/Bj20051939 (2006).

36 Lewis, E. R., Schwartz, S.E. Sea salt aerosol production: mechanisms, methods, measurements and models. Vol. 152 (American Geophysical Unit, 2004).

37 Daigle, C. C. et al. Ultrafine particle deposition in humans during rest and exercise. Inhal Toxicol 15, 539-552, doi:10.1080/08958370304468 (2003).

38 Henke, K. G., Sharratt, M., Pegelow, D. & Dempsey, J. A. Regulation of end-expiratory lung volume during exercise. J Appl Physiol (1985) 64, 135-146, doi:10.1152/jappl.1988.64.1.135 (1988).

39 Stokes, M. D. et al. A Marine Aerosol Reference Tank system as a breaking wave analogue for the production of foam and sea-spray aerosols. Atmos Meas Tech 6, 1085-1094, doi:10.5194/amt-6-1085-2013 (2013).

40 Orellana, G. et al. Validation of a confirmatory method for lipophilic marine toxins in shellfish using UHPLC-HR-Orbitrap MS. Anal Bioanal Chem 406, 5303-5312, doi:10.1007/s00216-014-7958–6 (2014).

41 Drust, B., Reilly, T. & Cable, N. T. Physiological responses to laboratory-based soccer-specific intermittent and continuous exercise. J Sport Sci 18, 885-892, doi:Doi 10.1080/026404100750017814 (2000).

42 Clarenbach, C. F., Senn, O., Brack, T., Kohler, M. & Bloch, K. E. Monitoring of ventilation during exercise by a portable respiratory inductive plethysmograph. Chest 128, 1282-1290, doi:DOI 10.1378/chest.128.3.1282 (2005).

43 Martin, M. Cutadapt removes adapter sequences from high-throughput sequencing reads. EMBnet.journal 17, doi:http://dx.doi.org/10.14806/ej.17.1.200 (2011).

44 Dobin, A. et al. STAR: ultrafast universal RNA-seq aligner. Bioinformatics 29, 15-21, doi:10.1093/bioinformatics/bts635 (2013).

45 Li, B. & Dewey, C. N. RSEM: accurate transcript quantification from RNA-Seq data with or without a reference genome. Bmc Bioinformatics 12, doi:Artn 32310.1186/1471-2105-12-323 (2011).

46 Robinson, M. D., McCarthy, D. J. & Smyth, G. K. edgeR: a Bioconductor package for differential expression analysis of digital gene expression data. Bioinformatics 26, 139-140, doi:10.1093/bioinformatics/btp616 (2010).

47 Nueda, M. J., Tarazona, S. & Conesa, A. Next maSigPro: updating maSigPro bioconductor package for RNA-seq time series. Bioinformatics 30, 2598-2602, doi:10.1093/bioinformatics/btu333 (2014).

